# Surface sediment samples from early age of seafloor exploration can provide a late 19th century baseline of the marine environment

**DOI:** 10.1101/419770

**Authors:** Marina C. Rillo, Michal Kucera, Thomas H. G. Ezard, C. Giles Miller

**Author notes:** Correspondence: Marina C. Rillo, Ocean and Earth Science, National Oceanography Centre Southampton, University of Southampton Waterfront Campus Southampton SO143ZH, UK,; C. Giles Miller, Department of Earth Sciences, Natural History Museum, Cromwell Road, London SW7 5BD, UK.

## Abstract

Ocean-floor sediment samples collected up to 150 years ago represent an important historical archive to benchmark global changes in the seafloor environment, such as species’ range shifts and invasions and pollution trends. Such benchmarking requires that the historical sediment samples represent the state of the environment at or shortly before the time of collection. However, early oceanographic expeditions sampled the ocean floor using devices like the sounding tube or a dredge, which potentially disturb the sediment surface and recover a mix of Holocene (surface) and deeper, Pleistocene sediments. Here we use climate-sensitive microfossils as a fast biometric method to assess if historical seafloor samples contain a mixture of modern and glacial sediments. Our assessment is based on comparing the composition of planktonic foraminifera (PF) assemblages in historical samples with Holocene and Last Glacial Maximum (LGM) global reference datasets. We show that eight out of the nine historical samples contain PF assemblages more similar to the Holocene than to the LGM PF assemblages, but the comparisons are only significant when there is a high local species’ temporal turnover (from the LGM to the Holocene). When analysing temporal turnover globally, we show that upwelling and temperate regions had greatest species turnover, which are areas where our methodology would be most diagnostic. Our results suggest that sediment samples from historical collections can provide a baseline of the state of marine ecosystems in the late 19th century, and thus be used to assess ocean global change trends.

## 1 INTRODUCTION

Late nineteenth and early twentieth century oceanographic expeditions set out to explore the vast and then widely unknown deep ocean. The voyage of the HMS *Challenger* is a notable example. As she sailed around the globe between 1872−76, researchers mapped for the first time the shape of the ocean basins and described over 4,500 new species of marine life (Manten, 1972). These early expeditions have important historical significance, as they mark the beginning of modern oceanography and stimulated further ocean exploration (Wüst, 1964).

From a scientific perspective, the observations and material acquired by these historical expeditions have great potential for global change research (Lister and Group, 2011; Johnson et al., 2011), as they provide a pre-1900 baseline of the marine environment (e.g., Roemmich et al. 2012; Gleckler et al. 2016). Yet historical seafloor sediment samples remain largely underutilised, because early seafloor sampling techniques involved collecting surficial sediments with instruments like the sounding tube, dredge or even the anchor (Thomson and Murray, 1891). All these instruments can penetrate below the surface and disturb the top layer of the sediment. As a result, such historical sediment samples might contain surface (Holocene) sediments mixed with deeper, glacial material (Hayward and Kawagata, 2005), potentially hindering their use as a historical baseline of the modern marine environment. Coring techniques provide more accurate sediment chronology (e.g., Röhl et al. 2000); however, historical samples represent the seafloor environment as much as 50 years earlier than the earliest core samples collected (Wüst, 1964), and thus contain sediments without any objects deposited after 1900. These uncontaminated historical samples can be useful for chemical analyses of the seafloor (e.g., pollution trends; Dekov et al. 2010), single-specimen analysis (e.g., Reichart et al. 2003; Wit et al. 2010) and investigations of species range shifts and invasions in the past century (e.g., Hoeksema et al. 2011). Therefore, it is important to assess the degree to which historical sediment samples represent Holocene or mixed-Pleistocene sediments.

One way to assess the degree of glacial mixing in the historical material would be to determine its absolute age using the radiocarbon dating technique, or to use glacial material proxies (e.g., Mg/Ca, oxygen isotopes). However, in cases where the extent of mixing is small, the exponentially decaying nature of the radiocarbon analysis can cause an ambiguous dating, and the isotopic analysis would require a large enough number of specimens to correctly represent the extent of the glacial mixing. Here we propose a complimentary method that uses planktonic foraminifera assemblage composition as a climate-sensitive fingerprint of the sediment age. Planktonic foraminifera (PF) are single-celled zooplankton that produce calcium carbonate shells and, upon death, accumulate in great numbers on the ocean floor (Hemleben et al., 1989). PF assemblage composition is sensitive to sea-surface temperature (Morey et al., 2005; Fenton et al., 2016) and its change between glacial and interglacial times has been used to determine the magnitude of glacial ocean cooling (MARGO Project Members, 2009).

In this brief report, we make use of the temperature sensitivity of PF and compare the composition of their assemblages in nine historical (*>* 100 years old) samples against reference PF assemblages from the Holocene (Siccha and Kucera, 2017) and the Last Glacial Maximum (Kucera et al., 2005a). We test whether it is possible to recover the extent of glacial mixing in historical seafloor sediment samples using PF assemblage composition. This new biometric method contributes to a more multidisciplinary approach to dating historical sediments.

## 2 MATERIAL AND METHODS

### 2.1 Historical samples

Historical samples were retrieved from the Ocean-Bottom Deposits (OBD) Collection held by The Natural History Museum in London. The OBD Collection holds about 40,000 historical samples from all the world’s oceans (https://doi.org/10.5519/0096416), including most of the sediment samples collected by the HMS *Challenger* and the British Royal Navy survey ships (Kempe and Buckley, 1987). The OBD samples are kept sealed in their original glass jars and tubes and are usually dry as the result of the long (over 100 years) storage. We selected nine samples collected between 1874 and 1905, chosen to cover different oceans, latitudes and historical marine expeditions (Table 1). Half of the amount available in the OBD containers was further split into two equal parts, leaving an archive sample and a sample to be processed. The sample processing consisted of weighing, wet washing over a 63*μ*m sieve and drying in a 60*°*C oven. The residues were further dry sieved over a 150*μ*m sieve and the coarser fraction was split with a microsplitter as many times as needed to produce a representative aliquot containing around 300 PF shells (see Al-Sabouni et al. 2007). All PF specimens in each of the nine final splits were picked, glued to a micropaleontology slide and identified under a stereomicroscope to species level, resulting in a total of 2,611 individuals belonging to 31 species (Table 1, Table S1).

**Table 1.** Information about the historical sediment samples from the Ocean Bottom Deposits (OBD) Collection at The Natural History Museum, London (NHMUK). Columns: NHMUK sample number; vessel that collected the sample (HMS stands for Her/His Majesty’s Ship); date that sample was collected; method used for sampling; ocean where sample was collected; latitude and longitude in decimal degrees; water depth in meters (transformed from fathoms); sampled mass before processing, in grams; number of splits to sample around 300 planktonic foraminifera (PF) individuals; number of PF individuals identified; number of PF species identified.

| Sample | Vessel HMS | Date | Method | Ocean | Lat | Long | Depth | m(g) | N(spl) | N(ind) | N(sp) |
| --- | --- | --- | --- | --- | --- | --- | --- | --- | --- | --- | --- |
| M.25 | *Challenger* | 21/02/1873 | Sounding | Atlantic | 24.33 | -24.47 | 5011 | 2.73 | 5 | 260 | 18 |
| M.192 | *Challenger* | 07/03/1874 | Sounding | Indian | -50.02 | 123.07 | 3976 | 0.19 | 5 | 318 | 7 |
| M.284 | *Challenger* | 11/03/1875 | Sounding | Pacific | -0.70 | 147.00 | 2012 | 1.98 | 7 | 331 | 16 |
| M.408 | *Challenger* | 21/03/1876 | Dredge | Atlantic | -21.25 | -14.03 | 3639 | 9.35 | 7 | 265 | 21 |
| M.3787 | *Egeria* | 31/10/1887 | Sounding | Indian | -19.57 | 64.63 | 2869 | 1.23 | 5 | 376 | 24 |
| M.4080 | *Egeria* | 26/08/1889 | Sounding | Pacific | -15.65 | -179.06 | 2579 | 2.42 | 8 | 300 | 13 |
| M.5246 | *Penguin* | 16/04/1891 | Sounding | Indian | -26.94 | 111.18 | 3676 | 1.49 | 5 | 279 | 18 |
| M.8780 | *Waterwitch* | 22/01/1895 | Sounding | Indian | -40.45 | 49.82 | 3780 | 1.51 | 6 | 177 | 11 |
| M.7487 | *Sealark* | 09/11/1905 | Sounding | Indian | -7.59 | 61.48 | 3758 | 2.86 | 8 | 305 | 18 |
| **Total** |  |  |  |  |  |  |  |  |  | 2,611 | 31 |

### 2.2 Holocene and Last Glacial Maximum data

We tested whether the composition of PF assemblages in the historical samples is more similar to assemblages of the Holocene (last 11,700 years, Walker et al. 2008) or the Last Glacial Maximum (LGM, 21,000 years ago, MARGO Project Members 2009). The Holocene census dataset (i.e., marine surface sediment samples taken by coring methods after 1945) was recently curated and published as the ForCenS dataset, comprising 4,205 assemblage counts from unique sites (Siccha and Kucera, 2017). Three LGM datasets from the MARGO project (Kucera et al., 2005a,b,c; Barrows and Juggins, 2004) were merged following the taxonomic standardisation of Siccha and Kucera (2017). This merged LGM final dataset included 1165 counts from 389 unique sites. Moreover, local estimates of open-ocean sedimentation rates are available for 156 samples in the LGM datasets, and were used to analyse the results.

The assemblage compositions of the nine historical samples were then compared to the samples from the geographically nearest site in the Holocene and LGM datasets (Fig. S1A). The distances between sites were calculated using the World Geodetic System of 1984 (WGS84, Hijmans 2015). We then compiled annual mean and standard deviation values of sea surface temperature (SST) from the World Ocean Atlas 2013 (WOA13, 0 meters depth, Locarnini et al. 2013) for each of the 27 sites (historical, Holocene and LGM) to evaluate whether the neighbouring sites are at similar SST ranges. Most neighbouring sites had similar SST values except the two most southern samples (Fig. S1B). The exceptional Holocene nearest sample neighbouring M.8780 was substituted by the fourth nearest neighbour, 66 km farther but more similar in SST (Fig. S1B). The mean distance between our nine sites and their nearest neighbour set was 253 km in the Holocene data and 415 km in the LGM data.

### 2.3 Compositional similarity

Assemblage similarity was expressed using the Morisita-Horn index (Morisita, 1959; Horn, 1966), which is an abundance-based overlap measure that preserves essential properties of similarity measurements (Jost et al., 2011). The Morisita-Horn calculates the compositional similarity by pairwise comparison of the relative abundance of each species, and is robust to under-sampling (i.e., rare species occurrence) (Jost et al., 2011). The index was calculated using the under-sampling bias correction (Chao et al., 2006), bootstrap confidence intervals based on 100 replicates and the R package *SpadeR* (version 0.1.1, Chao et al. 2016).

For each of the nine historical assemblages, we calculated the Morisita-Horn index three times: between (1) historical and neighbouring Holocene assemblages, (2) historical and neighbouring LGM assemblages and (3) neighbouring Holocene and LGM assemblages. This third comparison gives us a baseline index value of how much the PF assemblage composition changed locally since the LGM. If historical samples are representative of surface sediments, the similarity index calculated between historical and Holocene samples should be higher than between historical and LGM samples, with non-overlapping confidence intervals. However, if historical samples are a mixture of Holocene and LGM material, confidence intervals of the historical-Holocene and historical-LGM comparisons overlap. Confidence intervals might also overlap if the baseline index value calculated between the Holocene and the LGM reference samples is high. A high baseline value means that the local Holocene and LGM assemblages are similar and thus there is less statistical potential to detect whether a historical sediment is a surface or mixed-glacial sample. To understand where our biometric methodology would be most diagnostic, we compiled a world map of species turnover since the LGM, by calculating the Morisita-Horn index for each LGM sample and its nearest Holocene neighbour. Compositional similarity indexes were averaged per site. Distances between the Holocene sites and the 389 LGM sites were calculated using the WGS84, and averaged 52 km (median 1.5 km).

## 3 RESULTS AND DISCUSSION

In eight out of the nine samples, the compositional similarity was higher between the historical and Holocene assemblages than between historical and LGM assemblages (Fig. 1A). However, non-overlapping confidence intervals were only present in two samples (M.25 and M.5246), inferring a Holocene age for these historical sediments. These two samples also showed the lowest similarities in assemblage composition between neighbouring LGM and Holocene assemblages (Fig. 1A, grey dots), meaning that there was a greater change in PF assemblage composition since the LGM in these two locations. The main differences in species compositions were the high relative abundance of *Globigerina bulloides* in the LGM neighbouring sample of M.25, and of *Globoconella inflata* in the LGM neighbouring sample of M.5246 (Tables S3,S5).

**Figure 1.**
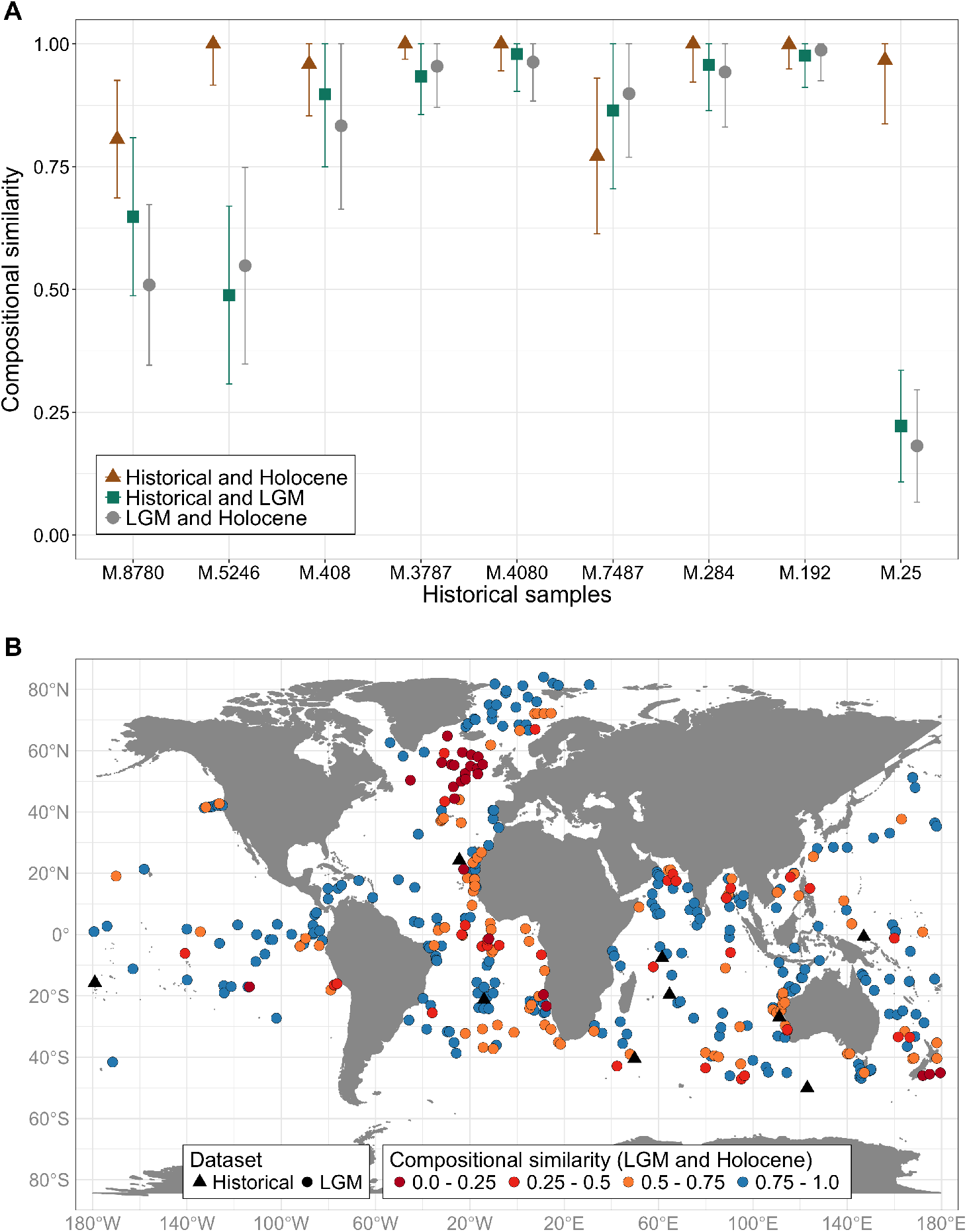
**(A)** Compositional similarity (Morisita-Horn index) between planktonic foraminiferal assemblages from historical sediment samples and assemblages from surface sediments (Holocene, brown triangle) and from the Last Glacial Maximum (LGM, green squares); and between Holocene and LGM assemblages (grey dots, baseline value of local temporal turnover). 0 means that the two assemblages share no species; 1 means that the same species were present in both samples at statistically indistinguishable proportions. Lines represent confidence intervals based on 100 bootstrap replicates. The x-axis shows the historical sample number. **(B)** Black triangles: historical samples (nine in total). Coloured dots: LGM samples from 389 sites worldwide. The colours represent the Morisita-Horn similarity index between the LGM sample and its neighbouring Holocene sample (i.e., temporal turnover). Red to orange dots indicate low similarity (i.e., high species turnover), whereas blue dots indicate similar Holocene and LGM planktonic foraminifera assemblages.

Compositional similarity index between historical and Holocene samples was always above 0.75, reaching maximum similarity in five samples (Fig. 1A). In six samples, the confidence intervals of the three comparisons overlapped, and all the similarity indexes were above 0.75. Since LGM and Holocene PF assemblages showed higher similarity at these six sites, our biometric test is less diagnostic. The historical sample M.8780 showed no overlap between historical-Holocene and Holocene-LGM, but the historical-LGM comparison overlapped with both comparisons, suggesting that either this sample has a mix of Holocene and glacial material, or the SST differences among these neighbouring samples prevents appropriate compositional comparisons (Fig. S1B). M.8780 had more *Neogloboquadrina pachyderma* than both the Holocene and LGM neighbours, and had *G. bulloides* abundances above 20%, similar to the LGM neighbour (Table S3). Moreover, differences in SST among the neighbouring samples of M.192 (Fig. S1B) did not seem to influence their compositional similarities (Fig. 1A). Finally, the historical sample M.7487 was the only one that showed higher similarity with LGM than Holocene assemblages, suggesting sediment mixing. The higher relative abundances of *Trilobatus sacculifer*, *N. incompta* and *Globigerinita glutinata* were responsible for this pattern (Table S5).

The global comparison between the Holocene and the LGM reference datasets shows the magnitude of PF assemblages turnover since the LGM (i.e., temporal beta-diversity, Fig. 1B). In general, upwelling (eastern boundary currents and equatorial regions) and temperate sites had greatest species turnover. Our methodology would be most diagnostic in these settings. Open-ocean sedimentation rates available for 156 sites averaged 6.8 centimetres per thousand of years (cm/ky). Therefore, historical sampling devices would have had to penetrate on average 142.8 cm (6.8 cm/ky times 21 ky) into the sediment to contaminate the surface seafloor sample with glacial material. Nevertheless, comparing sedimentation rates to the compositional similarity between LGM samples and their Holocene neighbours reveals that the greatest temporal turnover in species composition happened at sites of lower sedimentation rates (Fig. S2), where sediment mixing during historical sampling would be most likely. Furthermore, six LGM neighbours of our nine historical samples had local estimates of sedimentation rate, which varied from 1.0 to 3.4 cm/ky (mean 1.8 cm/ky, Tables S3,S4,S5). Thus, considering our historical samples only, depths of 21 – 71.4 cm into the sediment would have already reached glacial age material. Historical ocean-floor sampling methods potentially disturbed the surface and led to recovery of a mix of Holocene and deeper sediments, especially at sites of lower sedimentation rates. Our biometric method would be most useful at these sites with similar sedimentation settings, which also show the greatest temporal turnover in species composition (Fig. S2).

## 4 CONCLUSION

Our results indicate that historical ocean-floor sediment samples (collected more than 100 years ago) can represent surface (Holocene) sediments, despite the use of technology not designed to preserve undisturbed sediments. We show that the temporal turnover in species occurrence since the LGM varies in space making our biometric method particularly suitable for upwelling and temperate areas with low sedimentation rates. The new method allows a non-destructive preliminary assessment of glacial contamination of historical samples. Independent proxy records (e.g., Mg/Ca) and/or radiocarbon dating would be valuable to validate the success of our technique. Ideally, the biometric approach would compliment chemical-based techniques to date historical sediments, aiming for more robust results as multiple alternative lines of evidence are presented. Our results highlight the scientific potential of historical seafloor sediment collections (Table S2). As human activities increasingly modify the marine environment, these historical collections contain important information on the pre-1900 state of marine ecosystems.

## AUTHOR CONTRIBUTIONS

All authors designed the research question. MR processed the historical sediments, with input from GM. MR and MK identified all specimens. MR designed and performed the analysis, with input from MK and TE. MR wrote the initial draft, and all authors reviewed and edited the final manuscript.

## ACKNOWLEDGEMENTS

We are very grateful to Daniel Latorre for comments that improved the manuscript, to the MARGO Project members and to the ForCenS database authors for making their data openly available, to Dr Epi Vaccaro who made material from the Natural History Museum Ocean Bottom Deposits Collection available and to Andy Purvis for helping sample the museum collection.

## DATA AVAILABILITY STATEMENT

Upon acceptance of the manuscript, the datasets used and generated for this study will be available online at the NHM Data Portal (with DOI number), as well as the R Code used for the analysis and figures.

## SUPPLEMENTAL DATA

**Figure S1.**
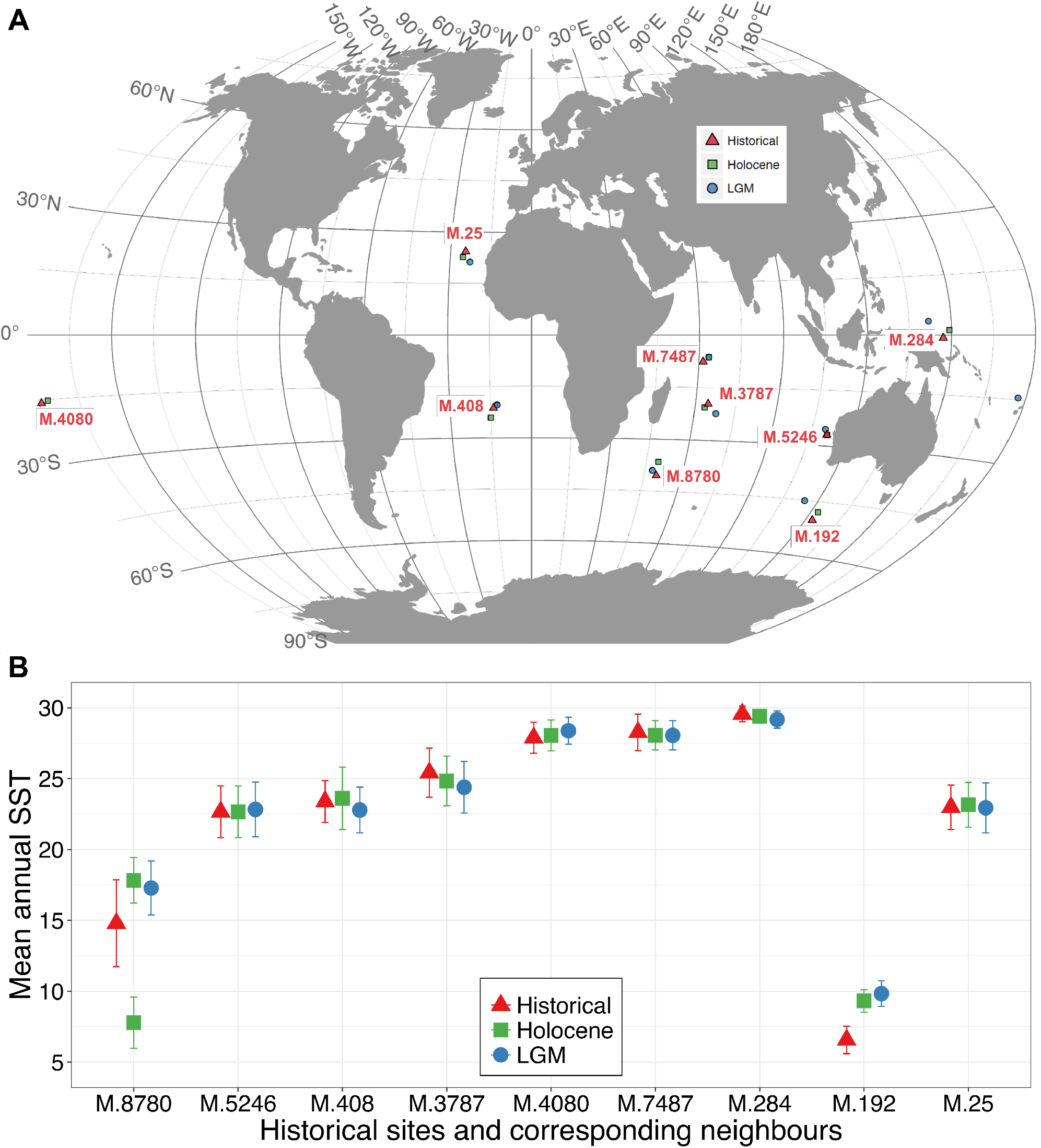
(A) Map of historical sample sites (red triangles) and corresponding neighbouring sites of the Holocene (green squares, ForCenS database) and Last Glacial Maximum (blue dots, LGM, MARGO datasets). Distances between each historical site and its corresponding Holocene and LGM neighbours can be seen in Tables S3,S4,S5. (B) Mean annual sea surface temperature (SST, in degree Celsius) and annual standard deviations (bars) at each sample site (WOA13 data). Note that historical sample M.8780 has two values for the Holocene neighbour. The nearest Holocene neighbour (SST 7.8*°*C, distance of 374 km) was substituted by the fourth nearest neighbour, which had more similar SST values (17.8*°*C, distance of 440 km, see map above).

**Figure S2.**
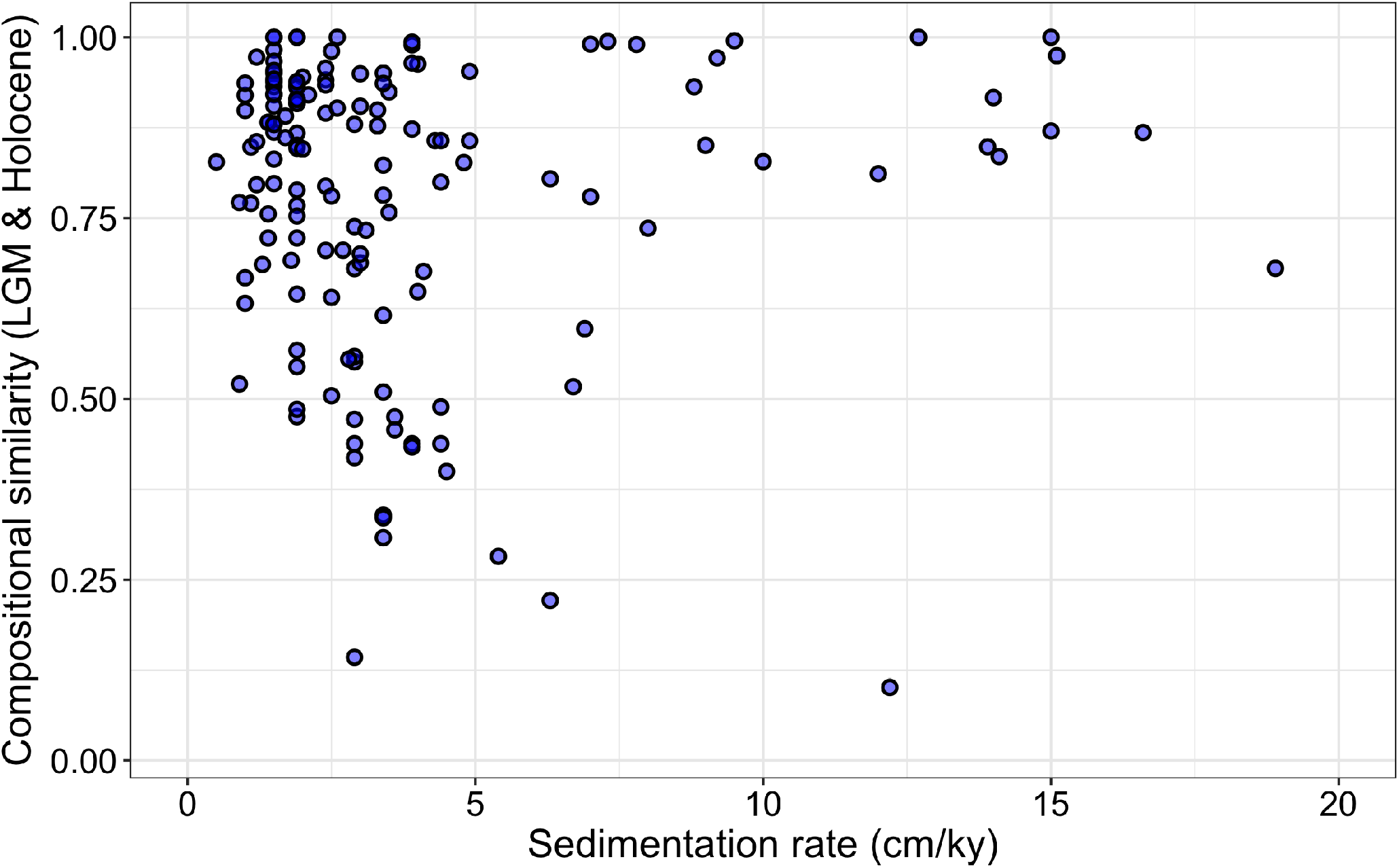
Compositional similarity (Morisita-Horn index) between planktonic foraminifera assemblages in the LGM sample and its neighbouring Holocene sample, plotted against local estimates of open-ocean sedimentation rates (centimetres per thousand of years). Low similarity means a greater temporal turnover in local assemblage composition. In total there are 150 sedimentation rate estimates; six samples with sedimentation rate higher than 20 cm/ky and compositional similarity above 0.75 are not shown.

**Table S1.**
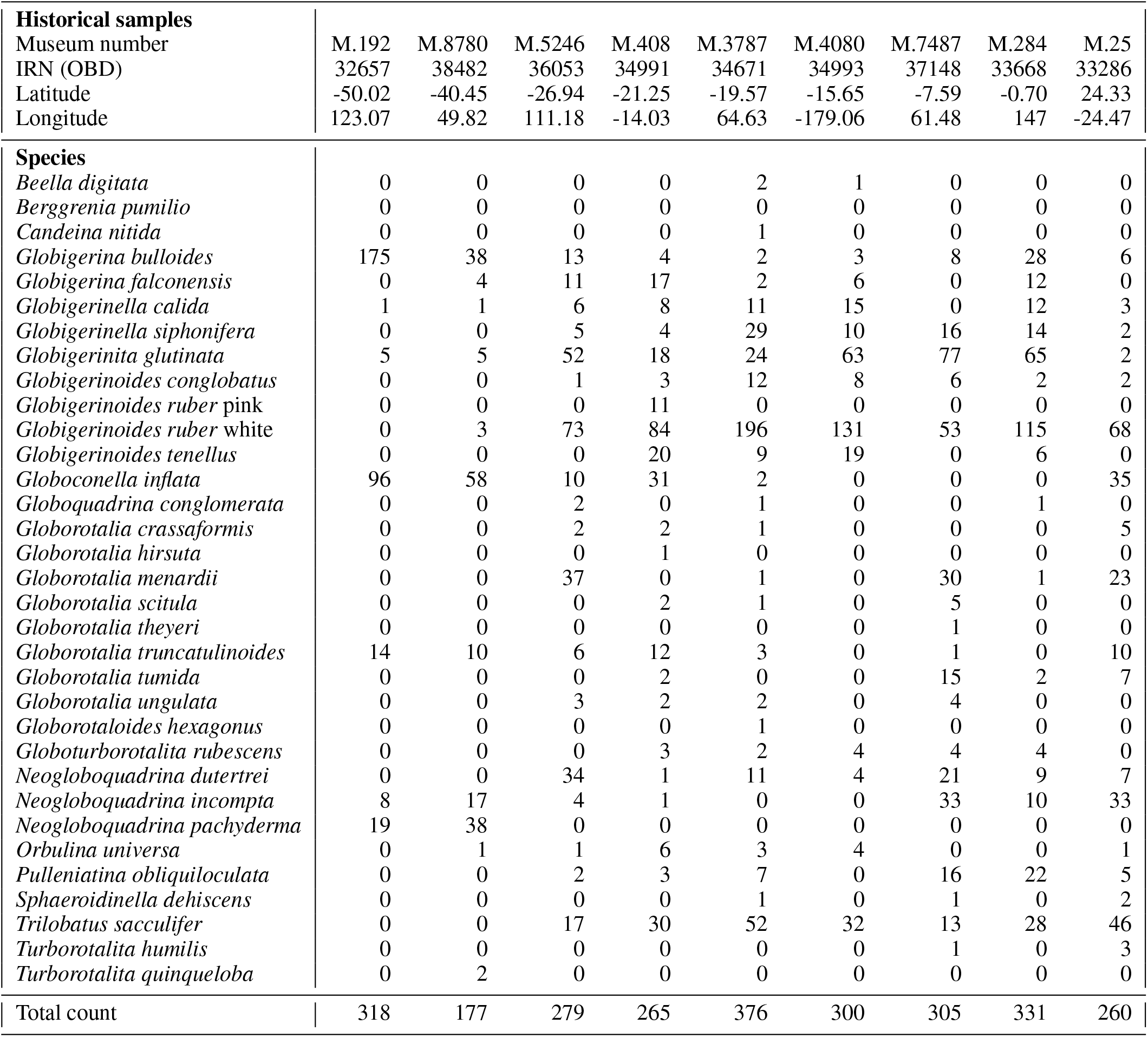
Planktonic foraminifera species counts of the nine historical samples (see Table 1). IRN (OBD) stands for the internal registration number of the Ocean-Bottom Deposits collection at the Natural History Museum in London. Individuals were identified to the species level by M. Rillo and M. Kucera together.

**Table S2.** Historical surface sediments collections

| Museum | Expedition | Spatial coverage | Period |
| --- | --- | --- | --- |
| Natural History Museum in London | *Challenger*<sup>1</sup> | Atlantic, Antarctic, South Indian, Pacific | 1872–76 |
| Muséum National d'histoire Naturelle in Paris | *Travailleur, Talisman*<sup>2</sup> | Atlantic, Mediterranean | 1880–83 |
| Museum für Naturkunde in Berlin | German Deep-Sea<sup>2</sup> (*Valdivia*) | Atlantic, Antarctic, Indian, Mediterranean | 1898–99 |

<sup>1</sup> https://www.hmschallenger.net/
<sup>2</sup> https://gallica.bnf.fr/ark:/12148/bpt6k65190274
<sup>3</sup> Keltie 1898; Sandeman 1900

**Table S3.**
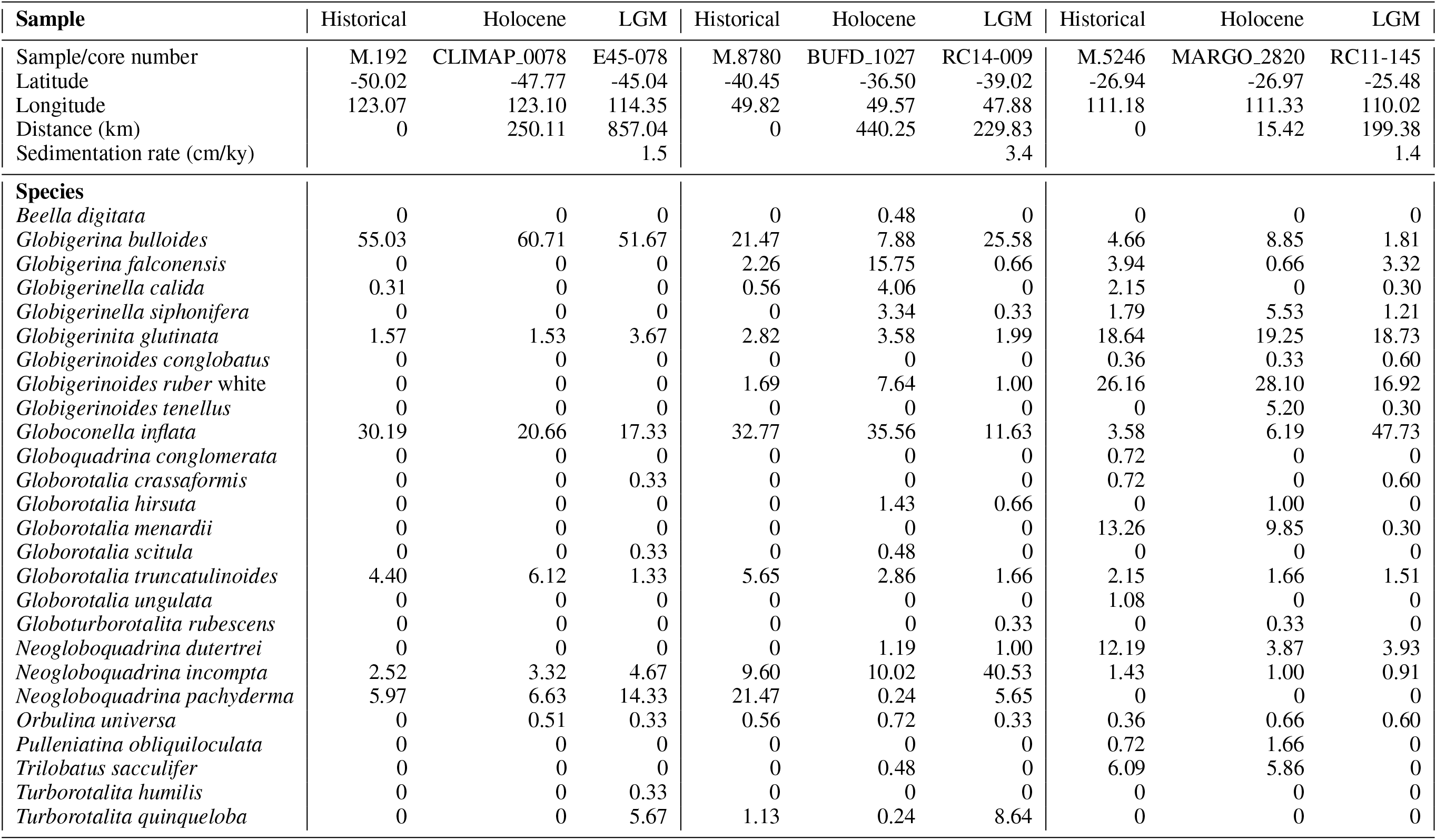
Relative abundance of planktonic foraminifera species in the historical samples M.192, M.8780 and M.5246, and their corresponding neighbouring samples in the Holocene and Last Glacial Maximum reference datasets. Species not shown had 0% abundance in all samples. Local sedimentation rate is given by the LGM datasets (see Methods) and expressed in centimetres per thousand of years.

**Table S4.**
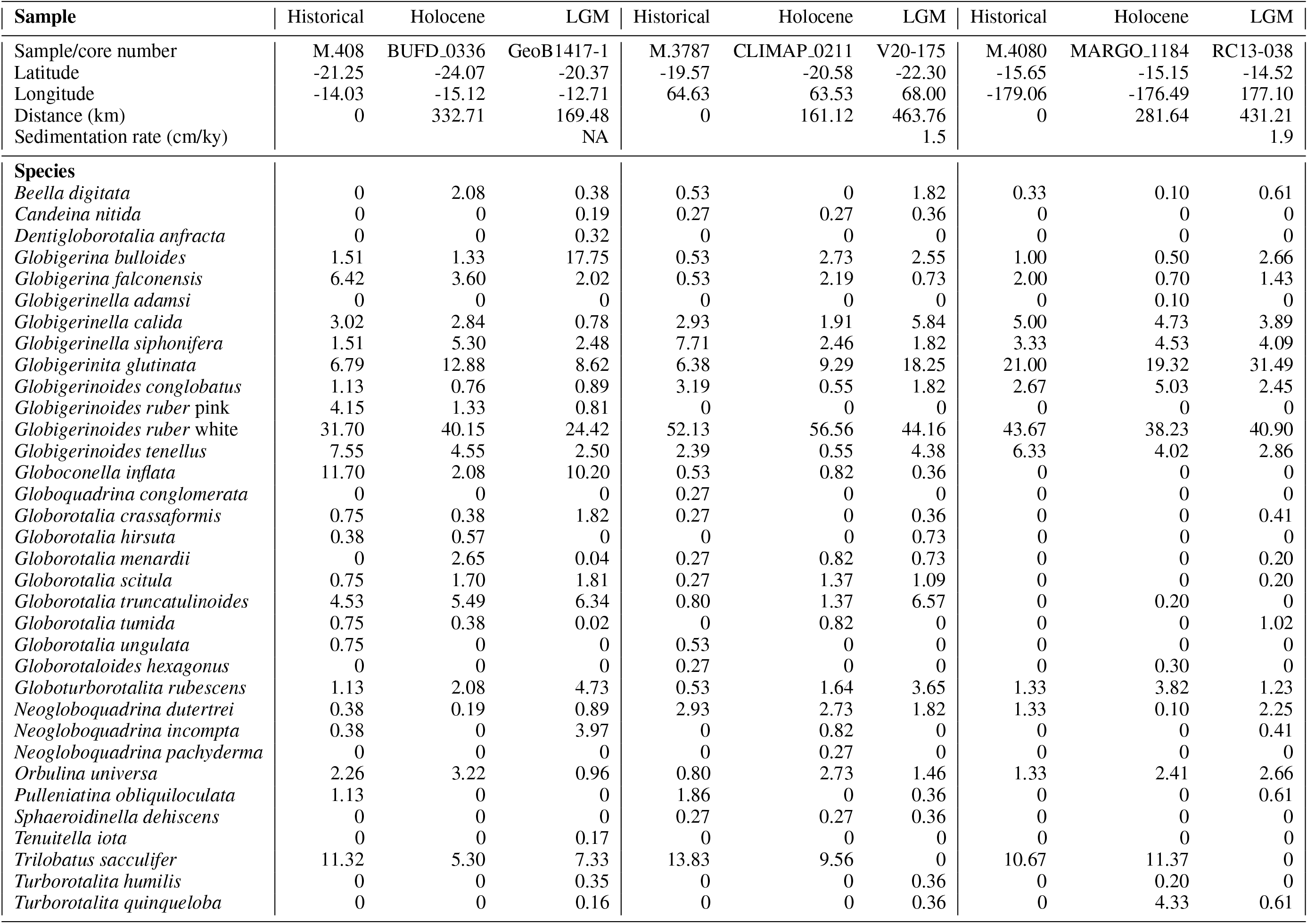
Relative abundance of planktonic foraminifera species in the historical samples M.408, M.3787 and M.4080, and their corresponding neighbouring samples in the Holocene and Last Glacial Maximum reference datasets. Species not shown had 0% abundance in all samples. Local sedimentation rate is given by the LGM datasets (see Methods) and expressed in centimetres per thousand of years.

**Table S5.**
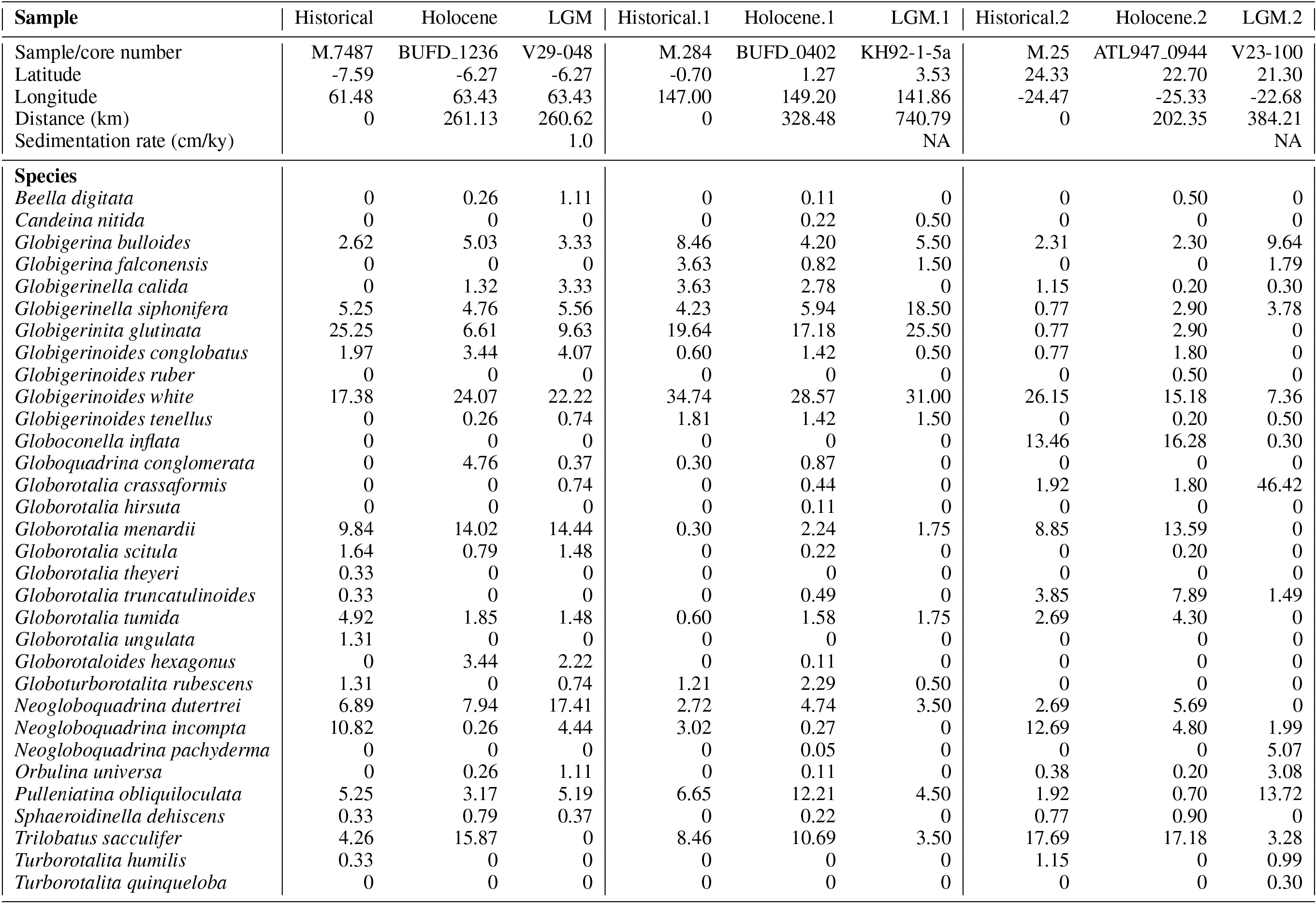
Relative abundance of planktonic foraminifera species in the historical samples M.7487, M.284 and M.25, and their corresponding neighbouring samples in the Holocene and Last Glacial Maximum reference datasets. Species not shown had 0% abundance in all samples. Local sedimentation rate is given by the LGM datasets (see Methods) and expressed in centimetres per thousand of years.

